# Mapping essential somatic hypermutations in a CD4-binding site bNAb informs HIV-1 vaccine design

**DOI:** 10.1101/2024.11.26.625503

**Authors:** Kim-Marie A. Dam, Harry B. Gristick, Yancheng E. Li, Zhi Yang, Priyanthi N.P. Gnanapragasam, Anthony P. West, Michael S. Seaman, Pamela J. Bjorkman

## Abstract

HIV-1 broadly-neutralizing antibodies (bNAbs) targeting the CD4-binding site (CD4bs) contain rare features that pose challenges to elicit these bNAbs through vaccination. The IOMA-class of CD4bs bNAbs includes fewer rare features and somatic hypermutations (SHMs) to achieve broad neutralization, thus presenting a potentially accessible pathway for vaccine-induced bNAb development. Here, we created a library of IOMA variants in which each SHM was individually reverted to the inferred germline (iGL) counterpart to investigate the roles of SHMs in conferring IOMA’s neutralization potency and breadth. Impacts on neutralization for each variant were evaluated, and this information was used to design minimally-mutated IOMA-class variants (IOMAmin) that incorporated the fewest SHMs required for achieving IOMA’s neutralization breadth. A cryo-EM structure of an IOMAmin variant bound to Env was used to further interpret characteristics of IOMA variants to elucidate how IOMA’s structural features correlate with its neutralization mechanism, informing the design of IOMA-targeting immunogens.

**Highlights:** IOMAmin variants with reduced SHMs retain neutralization potency and breadth.

Cryo-EM structure reveals IOMAmin preserves key Env interactions despite fewer SHMs.

IOMA variants with mutations in CDRH3 and CDRL1 fail to improve neutralization.

## Introduction

The discovery and extensive characterization of broadly neutralizing antibodies (bNAbs) against HIV-1 that potently neutralize a large fraction of circulating isolates have provided avenues to combat the ongoing HIV-1/AIDS pandemic, including the development of bNAbs for passive transfer and providing templates for vaccine design.^1–6^ bNAbs target several epitopes on Env, the sole viral glycoprotein on the surface of HIV-1, and 3D structures of HIV-1 Env-bNAb complexes have informed our understanding of bNAb features that confer diverse neutralization to enable structure-based immunogen design.^6–12^ Although substantial progress has been made on this front, an HIV-1 vaccine capable of eliciting bNAbs and providing robust and diverse protection has yet to be developed.

The binding site for CD4 (CD4bs), the HIV-1 host receptor, is an attractive target for HIV-1 immunogen design because bNAbs that recognize this epitope are among the most potent and broad.^6,12–17^ Many of these bNAbs share distinct V_H_ gene segments that encode particular features compatible with the CD4bs epitope and display impressive neutralization potencies.^12,14,18^ One class of CD4bs bNAbs that is V_H_1-2 gene segment-restricted includes two sub-classes: the well-studied VRC01-class,^9,13,14,16,18–21^ and the more recently described IOMA-class.^8,22^ Although these bNAb sub-classes share a V_H_ gene segment ontogeny, they have distinct sequence characteristics and binding mechanisms. The VRC01-class is defined by its rare, five-residue light chain complementary determining region (CDRL3) that is present in less than 1% of human antibody light chain genes.^13,18^ Other features include generally high levels of somatic hypermutation (SHM) and a CDRL1 with deletions or multiple glycine mutations necessary to accommodate the N276_gp120_ glycan.^13,18^ In contrast, the IOMA-class of CD4bs bNAbs are characterized by an 8-residue CDRL3, which is more commonly represented in the human B cell repertoire compared to VRC01-class five-residue CDRL3s.^8,22^ Additionally, IOMA-class Abs only require a single glycine mutation in CDRL1 to accommodate the N276_gp120_ glycan, a feature that is easier to achieve through vaccination than the VRC01-class CDRL1 changes. Finally, IOMA-class bNAbs have lower levels of SHM compared to the majority of bNAbs within the VRC01-class.^8,22^ High levels of SHM associated with HIV-1 CD4bs bNAbs is considered a hurdle in vaccine development^10,24^, as these mutations often require several years to accumulate during natural infection but are critical for achieving neutralization breadth.^24^

Current efforts to elicit CD4bs bNAbs have adopted a germline targeting approach, which seeks to create immunogens that specifically bind and activate inferred germline (iGL) forms of CD4bs bNAbs isolated from infected individuals.^25^ This method has been explored for the VRC01 and IOMA subclasses within the V_H_1-2 gene-restricted class of CD4bs bNAbs, since they have defined and well-characterized features.^26–31^ Although the VRC01-class is more broad and potent, the IOMA-class is thought to have features that are more feasible to elicit (i.e., a more common CDRL3 length, lower levels of SHM, a more easily achieved way to accommodate the N276_gp120_ glycan).^8,22,29^ Recent studies reporting the design and testing of IOMA-targeting sequential immunization strategies described eliciting CD4bs epitope–specific responses in wild-type animals and demonstrated broadly heterologous serum neutralization in knock-in and wildtype mice.^29^ Furthermore, IOMA-like monoclonal Abs isolated from these immunization studies developed mutations in CDRL1 that enabled accommodation of the N276_gp120_ glycan.^29^ Thus, directing germline focus towards IOMA-class antibody precursors presents a possible vaccine tactic for eliciting CD4bs bNAbs.

To further inform IOMA-targeting immunogen design, we evaluated IOMA’s structural features and identified characteristics that contribute to IOMA’s neutralization potency and breadth. We engineered a library of IOMA variants with reversions of SHMs to their iGL residues to identify which SHMs play a role in IOMA’s neutralization function versus which do not. Using this information, we created IOMAmin variants that included the minimal SHMs necessary to achieve the same neutralization potency and breadth as IOMA. Analysis of a 3.9 Å single-particle cryo-EM structure of an IOMAmin variant bound to Env showed that, despite containing fewer SHMs, the structural interactions between the IOMAmin variant and Env resembled those observed in the previously-characterized structure of mature IOMA bound to Env,^8^ validating our hypothesis that not all IOMA SHMs contribute to its ability to recognize and neutralize HIV-1.

Additionally, we explored whether mutations to IOMA’s CDRH3 and CDRL1 could improve neutralization function. The results illuminate IOMA’s mechanism of neutralization and also inform the design of IOMA-targeting immunogens and evaluation of potential IOMA-class bNAbs from natural infection or elicited by IOMA-targeting vaccine regimens.

## Results

### A fraction of SHMs contribute to IOMA’s neutralization potency and breadth

Although CD4bs bNAbs typically contain high levels of SHM,^13,14,18,21,22^ the design of minimally mutated CD4bs bNAbs and discovery of VRC01-class bNAbs with lower levels of SHM demonstrated that many SHMs are accessories of prolonged maturation during chronic infection and do not contribute to bNAb neutralization activity.^11,19,21,24^ Here, we sought to characterize the role of SHM in IOMA to understand how mutations influence neutralization and inform immunogen design efforts to elicit IOMA-class bNAbs. To identify SHMs in IOMA that contribute to its neutralization properties, we designed a panel of IOMA variants in which individual SHMs were reverted to their iGL counterparts and evaluated the effects of these substitutions on neutralization against a pseudovirus screening panel.

We designated two cohorts of IOMA SHMs: SHMs that contribute to the Ab:Env interface and interact with Env gp120 residues, termed internal face SHMs (inFACE), and SHMs that do not contribute to Ab:Env interactions, namely external face SHMs (exFACE). The inFACE and exFACE residues were distinguished by analyzing the BG505-IOMA (PDB 5T3X, 5T3Z) structures.^8^ SHM residues in the IOMA variable heavy (V_H_) and variable light (V_L_) domains containing an atom within 4.0 Å of a BG505 gp120 residue were considered inFACE residues and the remaining SHMs were exFACE residues. This analysis assigned 9 V_H_/9 V_L_ exFACE residues and 13 V_H_/7 V_L_ inFACE residues (Figures 1A, S1A).

**Figure 1:**
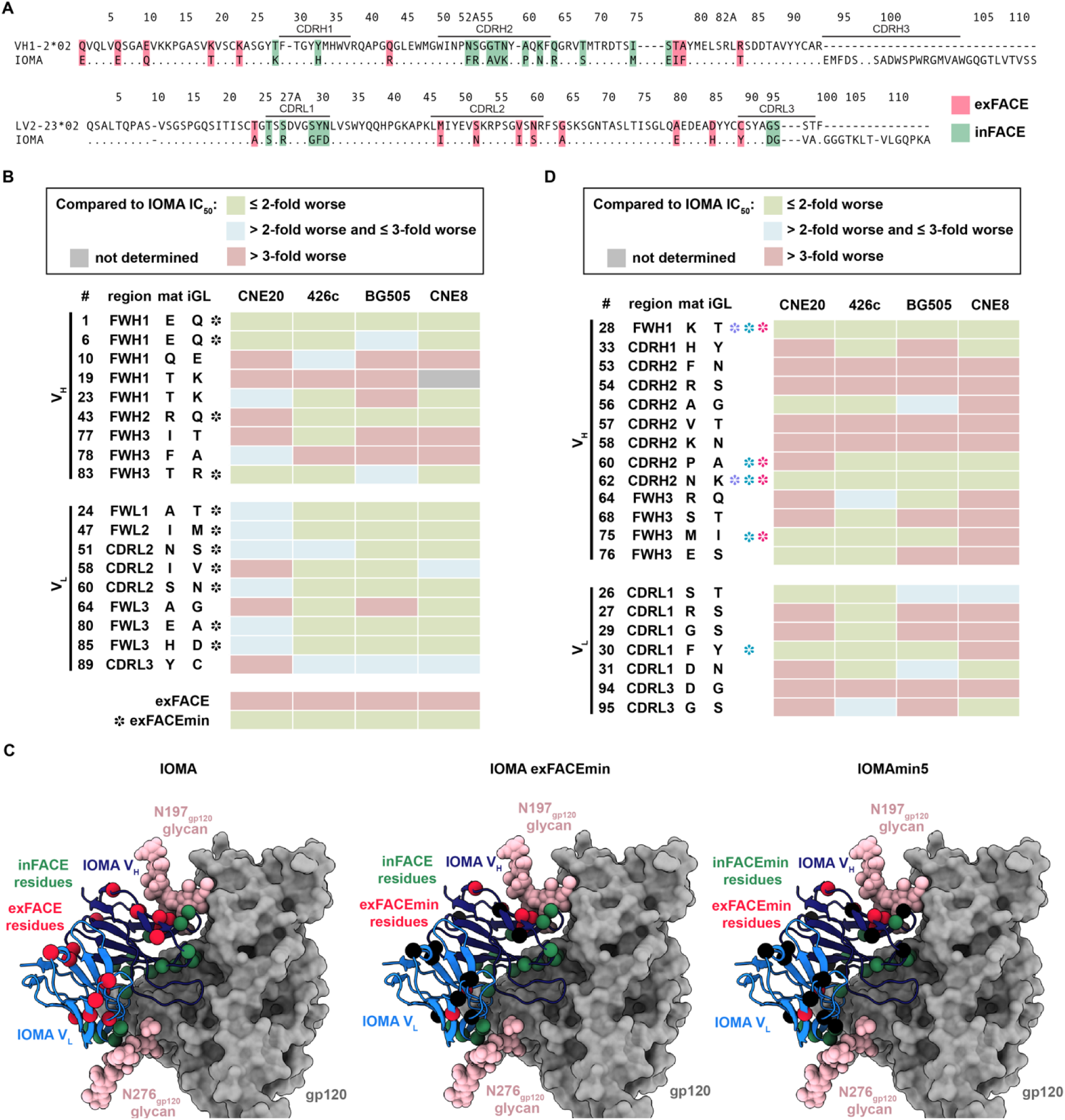
Screening SHMs that contribute to IOMA neutralization function. (A) Sequence alignment of IOMA iGL and mature V_H_/V_L_ sequences with exFACE and inFACE SHM residues highlighted in red and green, respectively. (B) Neutralization screening of IOMA exFACE variants against CNE20, 426c, BG505, CNE8 strains. Black symbols (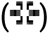) indicate exFACE mutations incorporated into the IOMAexFACEmin variant selected by the following criteria: IC_50_ values for each variant were ≤2-fold of IOMA’s IC_50_ against at least two strains and were >2-fold worse than mature IOMA against more than strain. (C) Structural representations of IOMA, IOMAexFACEmin, and IOMAmin5 Fabs bound to gp120 with exFACE and inFACE residues shown as red and green spheres, respectively (PDB 5T3X). Remaining SHMs are shown as black spheres. (D) Neutralization screening of IOMA inFACE variants against CNE20, 426c, BG505, CNE8 strains. Purple symbols (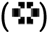) indicate exFACE mutations incorporated into the IOMAmin3 variant selected by the following criteria: IC_50_ values for each variant were within 2-fold of IOMA against all four strains. Teal symbols (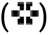) indicate exFACE mutations incorporated into the IOMAmin4 variant selected by the following criteria: IC_50_ values for each variant were within 2-fold of IOMA against at least three strains. Hot pink symbols (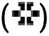) indicate exFACE mutations incorporated into the IOMAmin5 variant selected by the following criteria: IC_50_ values for each variant were within 2-fold of IOMA against at least three strains in the V_H_ and IC_50_ within 2-fold of IOMA against all four strains in the V_L_.

We first evaluated IOMA SHMs in the exFACE. Although these mutations were not involved in Ab:Env interactions, we hypothesized that some exFACE SHMs could contribute to maintaining IOMA’s structural integrity and thus indirectly contribute to neutralization potency and breadth. We created individual V_H_/V_L_ IOMA exFACE variants and tested neutralization against a small screening panel composed of pseudovirus strains that IOMA neutralizes with varying potencies (Figure 1B). Our panel included pseudoviruses for strains CNE20 (IOMA IC_50_: <0.1μg/mL), 426c (IOMA IC_50_: 0.1-0.99 μg/mL), BG505 (IOMA IC_50_: 1.0-9.9 μg/mL), and CNE8 (IOMA IC_50_: 10-50 μg/mL).8,32,33

Within V_H_, 4 of 9 exFACE variants (Q10E_HC_, T19K_HC_, I77T_HC_, F78A_HC_) resulted in > 3-fold decreases in IC_50_ compared to IOMA against 3 strains (Figure 1B). In the context of all SHMs in the exFACE, these 4 residues are in closest proximity to the N197_gp120_ N-glycan, suggesting these mutations may have evolved to stabilize glycan interactions (Figure S1B). In the V_L_, 7 of 9 exFACE variants resulted in comparable neutralization potencies compared to IOMA in the screening panel, suggesting exFACE SHMs in the LC play a lesser role in neutralization function compared to SHMs in the V_H_ exFACE (Figure 1B). Only 3 exFACE V_L_ variants (I58V_LC_, A64G_LC_, Y89C_LC_) exhibited >3-fold decreases in IC_50_ compared to IOMA against at least 1 strain (Figure 1B).

Based on these results, we designed an IOMAexFACEmin variant that incorporates the minimal number of SHMs in the exFACE required to maintain IOMA’s neutralization activity (Figure S1A). We applied the following criteria to select individual exFACE mutations: IC_50_ values for each variant must be ≤2-fold of IOMA’s IC_50_ against at least two strains and not be >2-fold worse in neutralization potency against IOMA for more than one strain (at least two green boxes and no more than one blue or red box) (Figure 1B,C). These mutations were then combined to create the IOMAexFACEmin variant. To validate our design of IOMAexFACEmin, we compared IOMA’s neutralization to IOMAexFACEmin and IOMAexFACE, a variant with all exFACE SHMs reverted to iGL residues, against the screening panel. We found that IOMAexFACEmin exhibited IC_50_ values within 2-fold of IOMA’s IC_50_ against all strains, whereas IOMAexFACE had IC_50_ values that were 3-fold greater than those of IOMA against all 4 strains (Figure 1B). These results are consistent with our hypothesis that a subset of somatically hypermutated exFACE residues contribute to IOMA’s neutralization activity without directly interacting with Env residues.

We applied the same methodology to IOMA inFACE residues (Figure 1D). Given that these residues interact with Env gp120, we expected that reversions of inFACE SHMs would adversely impact neutralization. Indeed, 8 of 12 V_H_ inFACE variants resulted in >3-fold increases in IC_50_ values compared to IOMA against at least 2 strains. 4 of these 8 variants (F53N_HC_, R54S_HC_, V57T_HC_, K58N_HC_) demonstrated weak neutralization against all strains (Figure 1D). Furthermore, 6 of 9 V_H_ inFACE variants within CDRH2 showed greatly impaired neutralization against at least 2 strains, indicating the importance of SHMs in this region during maturation of IOMA (Figure 1D). In the V_L_, 4 of 7 inFACE variants (R27S_LC_, G29S_LC_, D94G_LC_, G95S_LC_) resulted in >3-fold increases in IC_50_ compared to IOMA against at least 2 strains (Figure 1D). In one of these variants, G29_LC_ was reverted to the iGL Ser residue. A glycine in this position is hypothesized to facilitate the CDRL1 flexibility necessary to accommodate the N276_gp120_ glycan.^8^ Our results support the role of this substitution in IOMA’s neutralization function. Another CDRL1 inFACE variant, F30Y_LC_, did not result in an increase in IC_50_ compared to IOMA against three of the four screening strains, suggesting this mutation is not as important for N276_gp120_ glycan accommodation.

### IOMAmin variants show comparable neutralization potency and breadth to IOMA

Using results from the exFACE and inFACE single-site variant neutralization screens, we designed IOMAmin variants with the minimum numbers of SHMs required to maintain IOMA’s neutralization potency and breadth, first building upon the exFACEmin variant by incorporating inFACE mutations. We designed three variants based on the following criteria: IOMAmin3 included inFACE mutations with IC_50_ values within 2-fold of IOMA against all four strains (four green boxes), IOMAmin4 included inFACE mutations with IC_50_ values within 2-fold of IOMA against at least three strains (at least three green boxes), and IOMAmin5 inFACE mutations with IC_50_ values within 2-fold of IOMA against all four strains (four green boxes) in V_H_ and only exFACE mutations in V_L_ (Figures 1C, S1A).

The resulting IOMA variants contained a fraction of SHMs as IOMA, which has 22% somatically-mutated amino acid substitutions in V_H_ and 14% in V_L_. For IOMAexFACEmin, there were 18% and 8% somatically-mutated amino acid substitutions in V_H_ and V_L_, respectively (Figure S1C). The level of SHM substitutions for IOMAmin3, IOMAmin4, and IOMAmin5 were even further reduced to 16% V_H_ / 8% V_L_, 14% V_H_ / 7% V_L_, and 14% V_H_ / 8% V_L_, respectively (Figure S1C).

These variants contain the fewest SHM substitutions in the IOMA-class of CD4bs antibodies and similar levels of SHM substitutions to “minimal” VRC01-class antibodies such as minVRC01 and 12a21min^21^ (Figure S1C). We also investigated the effects of substitutions at these positions on non-specific binding using an in vitro polyreactivity assay for evaluating IgGs,^34^ finding that IOMA variants were not polyreactive as compared to highly polyreactive HIV-1 bNAbs such as 4E10 and 45-46m2^35,36^ (Figure S2D).

We next evaluated IOMAexFACEmin, IOMAmin3, IOMAmin4, and IOMAmin5 against a global 12-strain HIV-1 pseudovirus neutralization panel^37^ and six additional screening strains (Figure 2A). Notably, IOMAexFACEmin and IOMAmin5 exhibited comparable or lower geometric IC_50_ mean values against a global 12-strain panel^37^ compared to IOMA (4.5 and 5.3 μg/mL versus 6.5 μg/mL) while IOMAmin3 and IOMAmin4 showed comparable or higher geometric mean values against the 12-strain panel to IOMA (6.8 and 8.3 μg/mL versus 6.5 μg/mL). These trends in geometric mean IC_50_ values were consistent across all 18 strains (Figure 2A). IOMAmin4 exhibited the weakest geometric mean IC_50_s in both comparisons, suggesting a potential role of the F30Y_LC_ SHM in neutralization. Overall, these findings highlight the retention of neutralization activity for IOMAmin variants despite harboring fewer SHMs.

**Figure 2:**
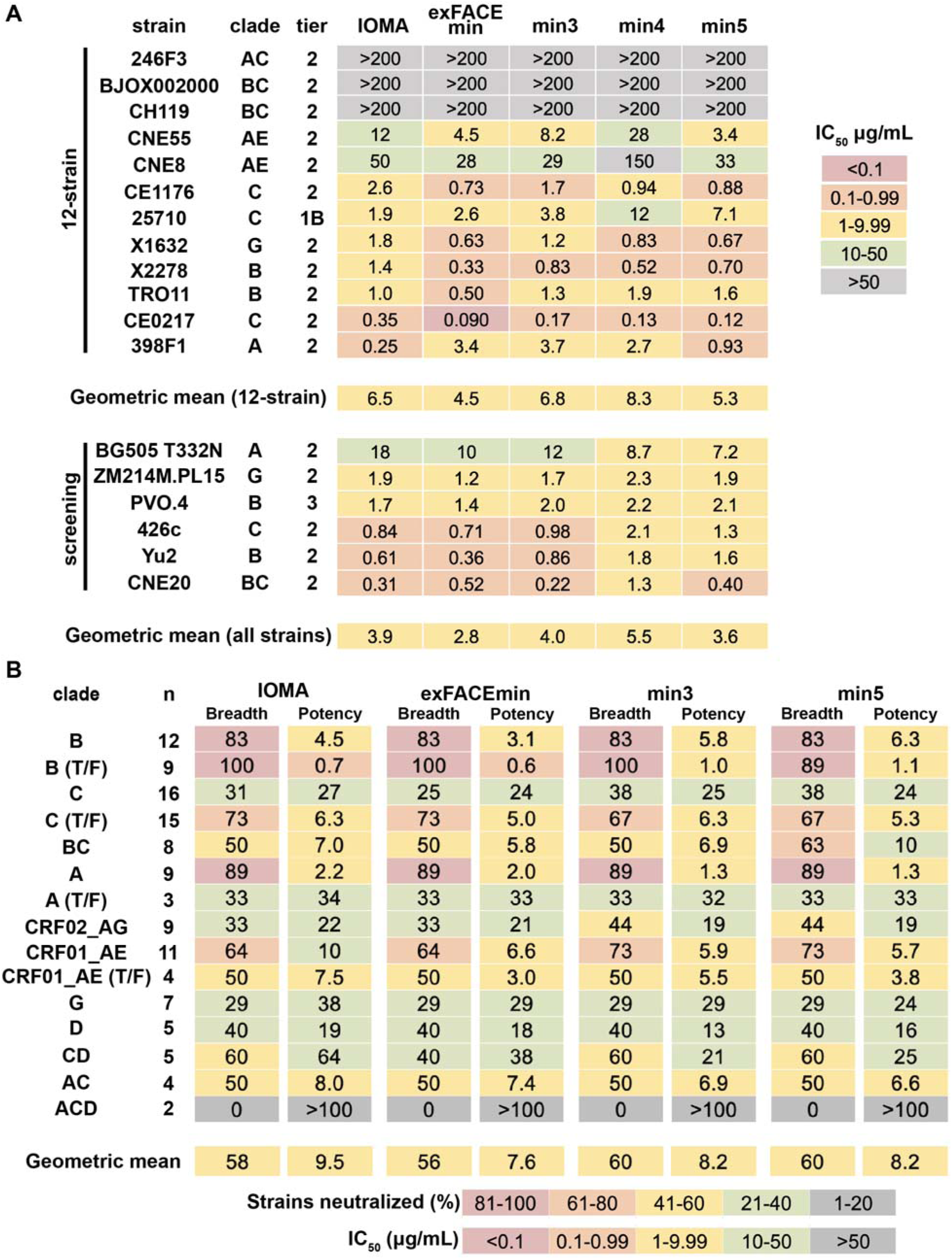
IOMAmin constructs show comparable neutralization profile to IOMA. (A) Neutralization IC_50_ values for IOMA and IOMAmin variants against the global 12-strain viral panel^37^ and six additional screening strains. (B) Overview of neutralization for IOMA and IOMAmin variants against a cross-clade 119-strain panel grouped into clades, where n is the number of strains per clade. Breadth is indicated by the percentage of neutralized strains for each clade (IC_50_ ≤ 50 µg/mL) and potency is indicated by geometric mean IC_50_.

We also evaluated IOMAexFACEmin, IOMAmin3, and IOMAmin5 against a cross-clade 119- strain pseudovirus panel to evaluate their potencies and breadth compared to mature IOMA, finding that all three variants showed improvements in potency compared to IOMA (Figure 2B). IOMAexFACEmin exhibited the highest potency with a geometric mean IC_50_ of 7.6 μg/mL, comparable to IOMA’s mean IC_50_ of 9.5 μg/mL. IOMAmin3 and IOMAmin5 also showed slight improvements in breadth, each neutralizing 60% of strains compared to IOMA’s 58%. These results emphasize the capacity of the IOMAmin variants to maintain, and even marginally enhance, their neutralization potencies and breadth despite having fewer SHMs.

### Structure of an IOMAmin5-Env complex shows different CDRH3 architecture from IOMA

To evaluate recognition of HIV-1 Env by IOMAmin5, we solved two single-particle cryo-EM structures of IOMAmin5 and 10-1074 Fabs bound to BG505 SOSIP Env, facilitating comparisons with a previously-described crystal structures of a IOMA and 10-1074 Fabs complexed with BG505^8^ (Figure 3A, S2A-B). Class I (3.9 Å resolution) revealed density for three IOMAmin5 and three 10-1074 Fabs bound to BG505, while class II (4.2 Å) displayed density for only two IOMAmin5 and three 10-1074 Fabs bound to BG505 (Figure S2C-F). Both classes exhibited C1 symmetry. Notably, despite approximate C3 symmetry in the class I structure, asymmetry was evident in the IOMAmin5 V_H_ CDRH3 in that each of the three IOMAmin5 CDRH3 regions adopted distinct disordered loops (Figure 3B). By contrast, the mature version of IOMA’s CDRH3, although also disordered, extends towards the CD4bs,^8^ unlike the CDRH3 loops observed for IOMAmin5 (Figure 3C). These differences may stem from the distinct sequences of IOMAmin5 and IOMA or from intrinsic disorder of CDRH3. Potential variations in the CDRH3s of mature IOMA bound to BG505 SOSIP could not be addressed because available structures of IOMA-BG505 complexes were derived from X-ray crystallographic analyses of complexes in which three-fold symmetry was imposed by crystal packing.^8^

**Figure 3:**
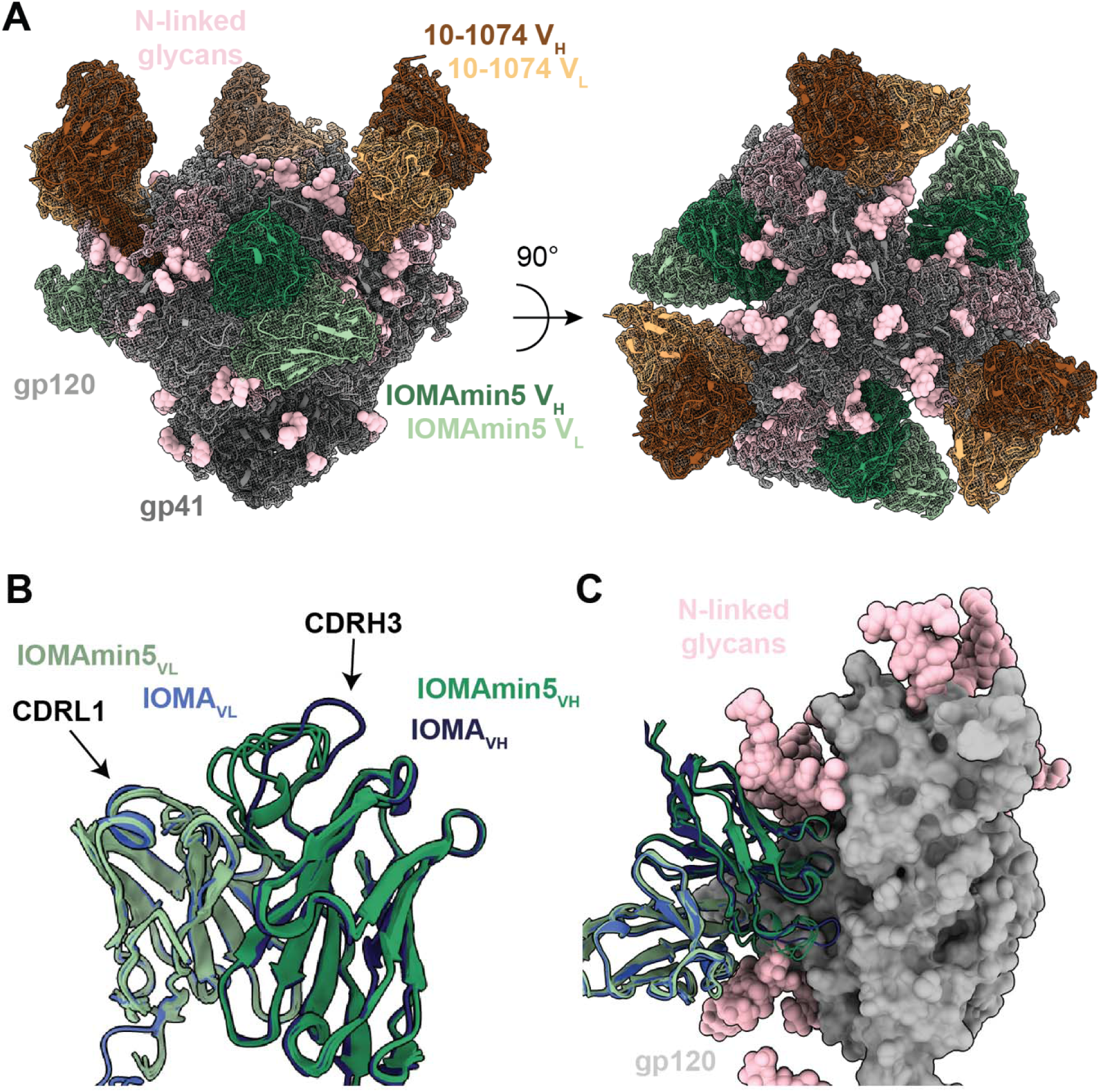
Single-particle cryo-EM structure of IOMAmin5-Env complex reveals altered CDRH3 architecture. (A) Side and top views of 3.9-Å single-particle cryo-EM density (mesh) and model (cartoon representation) of the IOMAmin5–BG505–10-1074 complex. N-linked glycans are represented as pink spheres. (B) Structural alignment of IOMAmin5 (this study) and mature IOMA (PDB 5T3X) Fabs (cartoon representation) with black arrows pointing to CDRL1 and CDRH3 loops. (C) Structural alignment of IOMAmin5 and IOMA V_H_-V_L_ domains (cartoon representations) bound to gp120 (surface representation) (PDB 5T3X).

A distinguishing characteristic of IOMA-class antibodies involves their mechanism for accommodating the N276_gp120_ N-glycan through CDRL1.^8^ In contrast to VRC01-class CD4bs bNAbs, which require a two- to six-residue deletion or selection of multiple glycines within CDRL1 to accommodate the N276_gp120_ glycan, IOMA-like bNAbs do not require such insertions or deletions.^8,13,14,18^ Instead, they rely on substitutions that enable a longer CDRL1 to provide the necessary flexibility to accommodate this glycan. The observation that IOMAmin5 adopts an identical CDRL1 conformation to that of mature IOMA (Figure 3B,C) suggests that sequence differences between IOMA and IOMAmin5 that are outside of CDRL1 (the CDRL1 sequences are identical) do not influence the CDRL1 conformation.

### IOMAmin5 exhibits CD4bs recognition comparable to that of IOMA

IOMAmin5 and IOMA buried comparable surface areas on BG505 gp120 (1020 and 1000 Å^2^, respectively); however, the distribution of buried surface area (BSA) on gp120 varied for IOMAmin5 and IOMA (Figure 4A, 4B). Unlike IOMA, IOMAmin5 showed no BSA in the gp120 inner domain, which includes the highly conserved K97_gp120_.^18^ Moreover, IOMA exhibited more than double the BSA in the gp120 exit loop compared to IOMAmin5, which had more concentrated BSA in the D and CD4 binding loops (Figure 4B). These distinctions in gp120 recognition may have contributed to slight variances in observed neutralization potencies (Figure 2).

**Figure 4:**
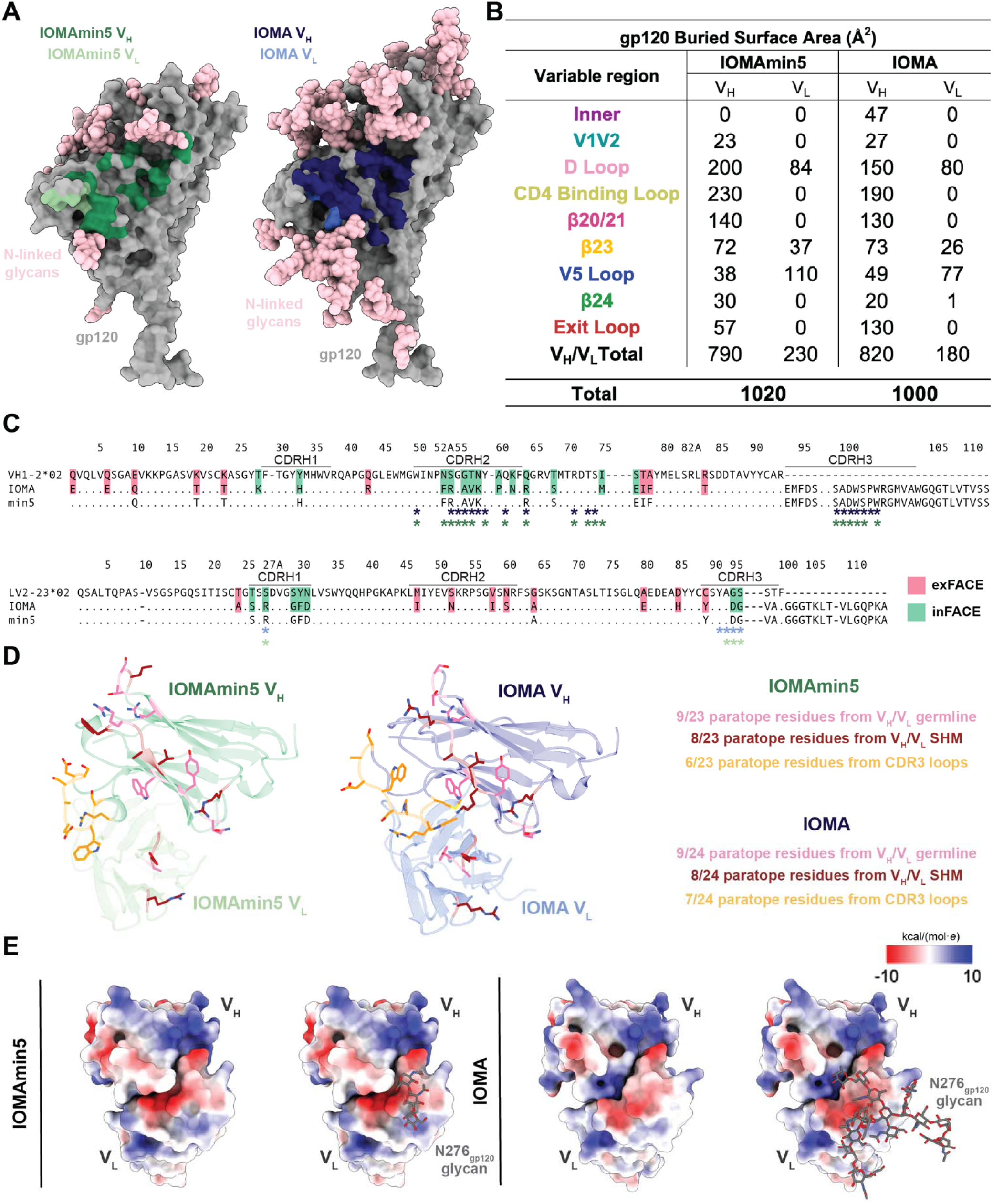
IOMAmin5 and mature IOMA exhibit similar gp120 CD4bs recognition. (A) Surface contacts made by IOMAmin5 V_H_ and V_L_ on BG505 gp120 (left) and surface contacts made by IOMA V_H_ and V_L_ on BG505 gp120 (right). (B) Summary of gp120 buried surface area (BSA) (Å^2^) calculations for IOMAmin5 and IOMA on regions of gp120: inner domain (inner), D loop, CD4bs loop, β20/21, β23, V5 loop, β24, and exit loop of the CD4bs. BSA calculations were conducted for gp120 protein components and did not include glycan interactions. (C) Sequence alignment of IOMA and IOMAmin5 (min5) constructs with iGL sequences. exFACE and inFACE SHM residues are highlighted in red and green, respectively. IOMA and IOMAmin5 V_H_/V_L_ residues that contribute to the paratope are labeled with an asterisk below the sequence alignment in blue (IOMA) and green (IOMAmin). (D) IOMA and IOMAmin5 paratope residues within CDR3 loops and from germline V genes or a SHM highlighted as sticks. (E) Surface representations of IOMAmin5 (left) and IOMA (right) V_H_-V_L_ domains with electrostatic potentials (kcal/(mol·*e*)) colored blue (positive electrostatic potential) to red (negative electrostatic potential) shown without and with superimposed coordinates for the N276_gp120_ glycan shown as sticks. Values were calculated using ChimeraX Coulombic Surface Coloring.^55^

A comparison of the paratopes of IOMAmin5 and IOMA revealed that residues engaged with the gp120 CD4bs were largely similar (Figure 4C). The most notable differences were observed in the CDRH2s, where IOMAmin5 exhibited interactions not seen in IOMA, utilizing F52_HC_, attributed to SHM, along with I75_HC_, which was one of the SHM reversions incorporated into the IOMAmin5 design. Both IOMAmin5 and IOMA utilized nearly 40% of the paratope surface for residues encoded by V_H_ and V_L_ germline genes and ∼30% for residues altered by SHM (Figure 4D). The remaining ∼30% of the paratope surfaces involved CDR3 loops (Figure 4D).

In addition to comparing the epitopes and paratopes of the IOMAmin5-BG505 and IOMA-BG505 structures, we evaluated potential electrostatic changes in the V_H_/V_L_-gp120 interfaces. Prior studies noted a shift towards a more positively-charged antigen combining site during the evolution of CD4bs-targeting bNAbs.^38^ This alteration was postulated to facilitate the accommodation of negatively-charged sialic acids on complex-type N-glycans in gp120.^38^ Although IOMAmin5 does not represent a naturally-occurring precursor of IOMA, we were interested in determining if the reversals of SHMs led to variations in the electrostatic potential of the antibody paratope. We found that regions of V_L_ near the N276_gp120_ glycan exhibited a somewhat more negative electrostatic potential for IOMAmin5 than for IOMA, suggesting a potential adaptation in IOMA to better accommodate the N276_gp120_ glycan, a complex-type N-glycan in BG505 SOSIP that includes negatively-charged sialic acids (Figure 4E).^39^

### Additional substitutions in CDRH3 and CDRL1 do not enhance neutralization by IOMA

In addition to optimizing IOMA’s level of SHM, we also engineered IOMA variants with modifications in CDRH3 and CDRL3 to investigate potential improvements in neutralization. The first variant, IOMA_HC-DDE_, included substitutions for CDRH3 residues S100_HC_, A100A_HC_, and D100B_HC_ to the negatively-charged DDE motif (Figure 5A), a motif that was first described for ACS103, an IOMA-class bNAb that was isolated from a person infected with HIV-1 who was characterized as an elite neutralizer.^22^ In addition, the DDE and similar motifs were identified in monoclonal antibodies isolated from an IOMA germline-targeting vaccination study in IOMA iGL knock-in mice.^29^ The latter study suggested that adoption of the DDE motif might have been driven by a well-conserved cluster of positively-charged residues present at the IOMA- contacting interface of the Envs utilized during the immunization regimen [K97_gp120_ (90% conserved), R476_gp120_ (R: 64% conserved; R/K: 98% conserved), and R480_gp120_ (99% conserved)].^29^ To assess the relevance of CDRL3 features in IOMA antibodies, we designed two IOMA variants with modifications in CDRL3. One variant, IOMA_5aa-CDRL3_, featured a 5-residue CDRL3 akin to the VRC01 class of bNAbs known for its broad and potent neutralization (Figure 5A).^13,14,18,20,38^ The other, IOMA_9aa-CDRL3_, featured a 9-residue CDRL3, a length that is nearly 8 times more prevalent in the human B cell repertoire than an 8-residue CDRL3 (Figure 5A).

**Figure 5:**
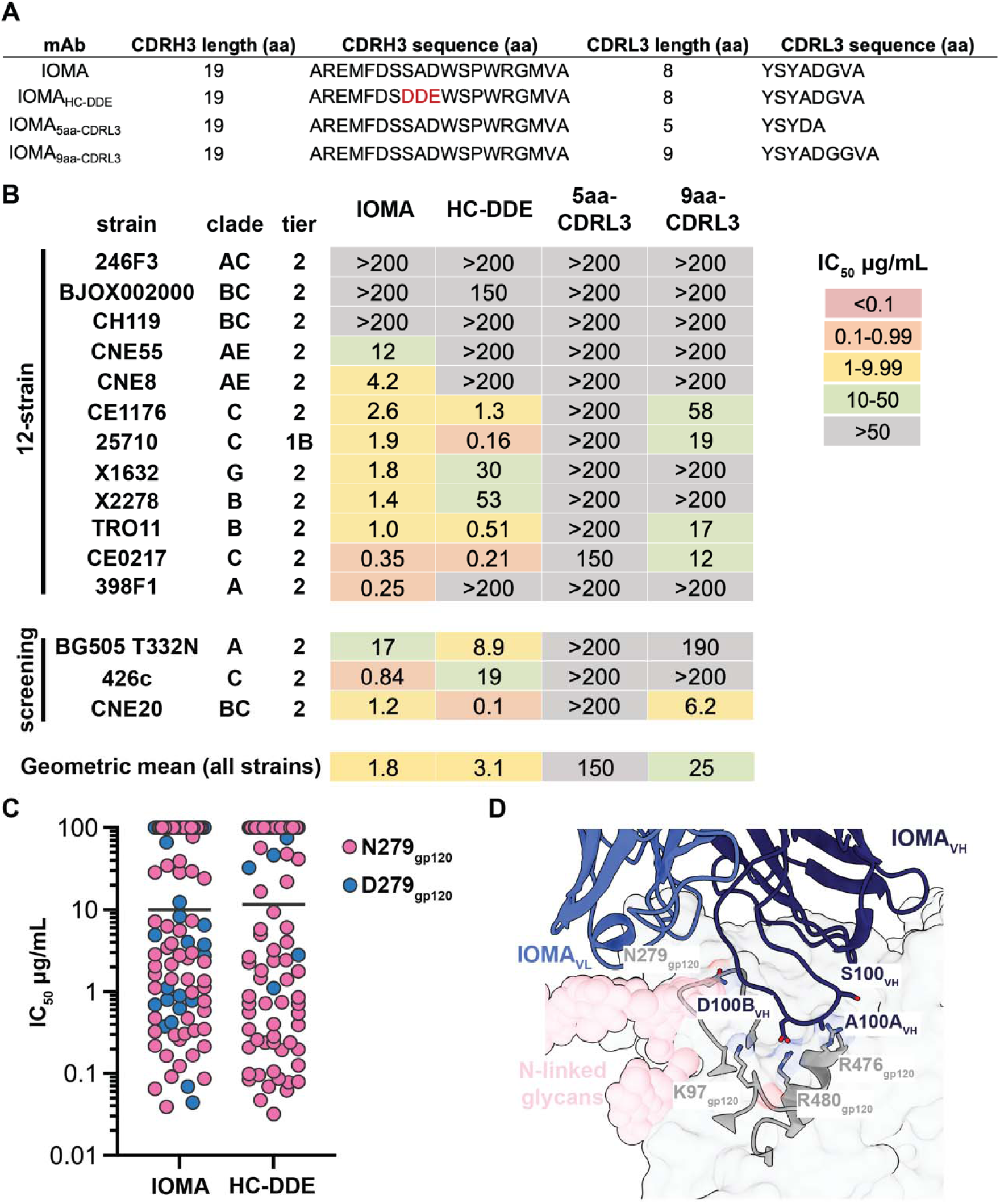
CDRH3 and CDRL1 mutations fail to improve IOMA neutralization. (A) Sequence alignment of CDRL1s and CDRL3s for IOMA and IOMA CDR variants. (B) Neutralization data for IOMA and IOMA CDR variants against a global 12-strain panel^37^ and three additional screening strains. (C) Neutralization IC_50_ profile of IOMA and IOMA_HC-DDE_ against a cross-clade 119-strain pseudovirus panel. Each symbol represents a unique HIV-1 isolate. Pink symbols represent isolates with N279_gp120_ and blue symbols represent isolates with D279_gp120._ (D) Interactions of IOMA CDRH3 with BG505 gp120 interface (PDB 5T3X).

We evaluated neutralization potencies of these variants in a global 12-strain HIV-1 pseudovirus neutralization panel^37^ plus three additional screening strains (Figure 5B). IOMA_5aa-CDRL3_ lost neutralization capabilities against all strains, while IOMA_9aa-CDRL3_ suffered a 10-fold increase in geometric mean IC_50_ compared to IOMA (25 versus 1.8 μg/mL). IOMA_HC-DDE_ had a slightly increased geometric mean IC_50_ compared to IOMA (3.1 versus 1.8 μg/mL), showing varied performance against different strains. To further explore this finding, we tested IOMA_HC-DDE_ against the cross-clade 119-strain pseudovirus panel and found that IOMA_HC-DDE_ is N279_gp120_- dependent (Figure 5C). This observation was unexpected, given that N279_gp120_ is not proximal to the well-conserved cluster of positively-charged residues in the CD4bs hypothesized to interact with the DDE motif (Figure 5D).^29^ This result could suggest that the DDE motif does not interact with the CD4bs in the hypothesized manner, possibly due to inherent flexibility in CDRH3, or that the DDE substitutions in this variant alter the conformation of CDRH3.

## Discussion

The identification and characterization of bNAbs against conserved HIV-1 Env epitopes provided a framework that led to engineering of immunogens to elicit bNAbs against particular epitopes and/or bNAbs of a specific class.^28,30,40–43^ Immunogen design to elicit bNAbs that bind the CD4bs is of particular interest since CD4bs bNAbs have been well-characterized and are among the most broad and potent anti-HIV-1 bNAbs.^18,25^ However, the CD4bs epitope involves a recessed pocket that is framed by bulky N-linked glycans, presenting steric challenges that require CD4bs bNAbs to adopt rare features, further challenging vaccine design efforts.18,24,25,44,45

In this study, we focused on the IOMA-class of CD4bs bNAbs, which represent a promising target due to their distinct characteristics: lower levels of SHM compared to most CD4bs bNAbs, an 8-amino acid CDRL3, and fewer mutations in CDRL1 to accommodate the N276_gp120_ glycan.^8,22,29^ We further characterized IOMA’s neutralization mechanism by evaluating the role of SHMs. We identified 5 of 9 V_H_ and 2 of 9 V_L_ exFACE SHMs that contribute to neutralization by IOMA but are not predicted to interact with Env gp120 residues. In the V_H_ domain, the proximity of these mutations to the N197_gp120_ glycan suggested they evolved to stabilize and/or accommodate this glycan. Furthermore, we found that almost half of all V_L_ SHMs were in the exFACE and appeared to play no role in neutralization by IOMA. This suggested that V_L_ accumulated “passenger” mutations, likely resulting from IOMA’s prolonged SHM process. For inFACE SHMs that were predicted to contact gp120 residues, we found that most mutations in V_H_ and V_L_ contribute to IOMA’s neutralization activity. In particular, inFACE SHMs in CDRH2 and in CDRL1, adjacent to the N276_gp120_ glycan when IOMA interacts with Env, proved to be necessary for IOMA’s potency and breadth, further suggesting that N-glycans in the gp120 CD4bs stimulated the evolution of IOMA’s SHMs.

With these results, we generated three IOMAmin variants that each included a fraction of the SHM substitutions found in mature IOMA but maintained its neutralization potency and breadth, suggesting that relatively few SHMs are needed for IOMA-class cross-neutralizing activity.

These results are useful for identifying sequences of IOMA-like antibodies derived from human patients and animal studies because they can guide and possibly predict specific mutations that are important for broad and potent neutralization. Furthermore, the identification of key IOMA- gp120 interactions can be used to inform germline-targeting immunogen design.

Our findings also demonstrated that the mutations we introduced into IOMA’s HC and LC CDR3 regions did not result in improved neutralization. For example, engineering of a VRC01-like five-residue CDRL3 or a more prevalent nine-residue CDRL3 both resulted in reduced neutralization potencies. These outcomes raise intriguing questions regarding the structural and functional consequences of these mutations. Further structural characterization of these variants is warranted to unravel the underlying mechanisms and explore the feasibility of optimizing CDRL3 sequences to enhance their neutralization potentials.

In addition, our investigation of the IOMA_HC-DDE_ variant revealed unexpected results. We hypothesized that the incorporation of the negatively-charged DDE motif in CDRH3^22,40^ would be advantageous, potentially enhancing neutralization by binding to a conserved cluster of positively charged residues at the interface with Env. However, contrary to this hypothesis, the incorporation of the CDRH3 DDE motif failed to augment neutralization capability compared to mature IOMA. Instead, the presence of the DDE motif skewed neutralization towards N279_gp120_- dependency, thereby impairing neutralization efficacy. This observation is particularly noteworthy considering that only half of Env strains contain asparagine at position 279_gp120_.^33^ Indeed, the selection of the DDE motif by the human antibody ACS103 during a naturally occurring HIV-1 infection demonstrated that these residues conferred an advantage in protecting from the circulating HIV-1 strains in that individual.^22^ However, the decreased breadth of ACS103 compared to IOMA^22^ suggests that the presence of the DDE motif does not increase breadth or potency across diverse HIV-1 strains, consistent with results presented here. The implications of this outcome underscore the complexity of antibody-Env interactions and highlight the need for a more comprehensive understanding of factors influencing neutralization breadth and potency.

Together, the identification of mutations in IOMA that confer bNAb neutralization activity reveal viral pressures that influenced IOMA’s development. Furthermore, we anticipate that this information can be used to not only engineer IOMA-targeting immunogens and evaluate potential IOMA-class bNAb sequences isolated from experimental vaccine regimens, but also to inform the design of improved CD4bs immunogens to elicit antibodies with increased breadth and potency across diverse HIV-1 strains.

### Experimental Procedures Design of IOMA variants

Sites for IOMAmin exFACE and inFACE mutations were determined by analyzing the interactions between BG505 and IOMA in the BG505-IOMA-10-1074 structures (PDB 5T3Z and 5T3X) in PyMol (Schrödinger LLC). IOMA residues that include atoms that were ≤4.0 Å from a BG505 residue were determined as inFACE residues and the remaining SHMs were defined as exFACE residues. Genes encoding IOMA exFACE and inFACE single-site variants and IOMA CDRL3 and DDE variants were generated using site-directed mutagenesis (Agilent QuikChange) starting with mature IOMA HC and LC genes, and IOMAexFACEmin and IOMAmin variants were generated using Gibson cloning (NEB Gibson Assembly).

### Protein expression and purification

Expression vectors encoding IgGs and Fabs were transfected using the transient Expi293 expression system (ThermoFisher), according to the manufacturer’s protocol.^7,46^ Expression vectors included IgG HC or Fab HC and LC genes. Fab HC expression vectors encoded a C-terminal 6x-His tag. Expressed proteins were isolated from cell supernatants from IgG and Fab transfections using protein A (GE Healthcare) and Ni^2+^-NTA (GE Healthcare) affinity chromatography for IgGs and Fabs, respectively. Subsequently, IgG and Fabs were purified over size exclusion chromatography (SEC) using a Superdex 200 10/300 column (GE Healthcare). Proper folding of variants was assessed by SEC profiles and expression yields.

BG505 SOSIP.664 Env constructs encoded SOSIP mutations including disulfide mutations 501C and 605C (SOS), I559P (IP), and the furin cleavage site mutated to six arginine residues (6R).^47^ BG505 SOSIP.664 Env expression vectors were transfected using the transient Expi293 expression system (ThermoFisher), according to the manufacturer’s protocol. Trimeric Env was separated from cell supernatants using PGT145 immunoaffinity chromatography and SEC using a Superose 6 10/300 column (GE Healthcare).^47,48^

### HIV-1 TZM.bl Neutralization Assays

Neutralization activities of IOMA-based IgGs were determined using a luciferase-based TZM.bl pseudovirus assay conducted using standard protocols.^37,49^ IC_50_ values were determined from independent replicates (n=2) analyzed using Antibody Database (v2.0)^33^ with 5-parameter curve fitting. Non-specific activity was determined by evaluating IgGs against murine leukemia virus (MuLV).

### Polyreactivity Assay

A baculovirus-based polyreactivity assay was performed using an established enzyme-linked immunosorbent assay (ELISA) method to detect nonspecific binding.^34^ Briefly, a solution containing 1% baculovirus particles in 100 mM sodium bicarbonate buffer (pH 9.6) was applied to the wells of a 384-well ELISA plate (Nunc Maxisorp) using a Tecan Freedom Evo liquid handling robot. Following overnight at 4°C incubation, plates were blocked for 1-hour at room temperature with phosphate-buffered saline (PBS) containing 0.5% bovine serum albumin.

Purified IgGs, diluted to 1 µg/ml in PBS with 0.5% bovine serum albumin, were added to the blocked assay plate and incubated for 3 hours at room temperature. Detection of bound IgG utilized a horseradish peroxidase–conjugated anti-human IgG (H&L) secondary antibody (GenScript), with luminescence signals measured at 425 nm with SuperSignal ELISA Femto Maximum Sensitivity Substrate (Thermo Fisher Scientific).

### Assembly of protein complexes and cryo-EM sample preparation

Protein complexes for cryo-EM were generated by combining purified IOMAmin5 and 10-1074 Fabs with a BG505 SOSIP.664 Env trimer in a 3:1 Fab:trimer molar ratio and incubating at 4°C overnight. Fab-Env complexes (3 μL) were applied to Quantifoil R2/2 400 mesh cryo-EM grids (Ted Pella) that were prepared by glow-discharging for 1 min at 20 mA using a PELCO easiGLOW (Ted Pella). Grids were blotted with Whatman No. 1 filter paper for 3 s at 100% humidity at room temperature and vitrified via plunge-freezing in liquid ethane using a Mark IV Vitrobot (ThermoFisher).

### Cryo-EM data collection

For the single-particle cryo-EM study of the IOMAmin5-BG505-10-1074 complex, data collection was performed using a Titan Krios transmission electron microscope operated at 300LkV. Movies were recorded with beam-image shift over a single exposure per hole in a 3-by-3 pattern of 2 μm holes. Movies were captured in super-resolution mode on a K3 camera (Gatan) equipped with a BioQuantum energy filter (Gatan) using a 20LeV slit width, yielding a pixel size of 0.4327LÅ•pixel^−1^. The defocus range was set between 1.0 and 3.0 μm.

Data processing was performed with RELION software.^50,51^ Movies were motion corrected using MotionCor2^52^ after binning, and GCTF^53^ was used for contrast transfer function (CTF) estimation. Micrographs with poor CTF fits or ice quality were excluded. Manual particle picking was conducted for a subset of particles, followed by reference-free 2D classification. Selected 2D class averages were used for automated particle picking using the RELION AutoPicking module.^50,51^ Resulting particles were subjected to several rounds of 2D and 3D classifications.

An initial model was generated using cryoSPARC with a subset of particles and used as a reference for 3D classification assuming C1 symmetry.

Two distinct classes were identified from 3D classification representing the IOMAmin5-BG505- 10-1074 complex: class I exhibited density corresponding to three IOMAmin5 and three 10-1074 Fabs bound to BG505; class II showed density for two IOMAmin5 and three 10-1074 Fabs bound to BG505. These classes were subsequently refined using 3D auto-refinement and underwent post-processing in RELION.^50,51^ Particle polishing, CTF refinement, and multiple iterations of 3D auto-refinement were conducted to enhance map quality. Final high-resolution maps were generated and resolutions were determined in RELION^50,51^ based on the gold-standard Fourier shell correlation (FSC) criterion at 0.143. FSCs were computed using the 3DFSC program.^54^

### Cryo-EM model building and refinement

Models were generated by fitting the coordinates of gp120 (PDB 5T3X), gp41 (PDB 5T3X), 10- 1074 Fab (PDB 5T3X) into cryo-EM density maps using UCSF ChimeraX.^55^ Initial models were refined using the Phenix program’s real-space refinement module.^56,57^ Subsequent updates to the sequence and additional manual adjustments were carried out using Coot software.^58^ The refinement process involved iterative cycles of automated refinement using Phenix^56,57^ and manual adjustments in Coot to produce the final models (Table S1).^58^

### Structural analyses

PyMol (Schrödinger LLC) and UCSF ChimeraX^55^ were used to prepare structure figures. PDBePISA was used to calculate BSAs using a 1.4 ALJ probe. BSA calculations for gp120 were for its protein components and did not include contributions from glycans. Defined interactions were assigned tentatively due to the relatively low resolution of complexes using the following criteria: hydrogen bonds were assigned for pairwise interactions <4.0 Å and with an A-D-H angle >90° and van der Waals interactions were assigned as distances of <4.0 Å between atoms. Electrostatic potentials were calculated using the Coulombic Surface Coloring module in UCSF ChimeraX.^55^

## Acknowledgements

We thank J. Vielmetter, A. Rorick, K. Storm, and the Protein Expression Center in the Beckman Institute at Caltech for expression assistance, and Z. Wu and J. Keeffe for help with polyreactivity assays. Electron microscopy was performed in the Caltech Cryo-EM Center with assistance from S. Chen. This work was supported by the National Institute of Allergy and Infectious Diseases (NIAID) Grant HIVRAD P01 AI100148 (P.J.B.) and NIH 1U54AI170856 (P.J.B.). The contents of this publication are solely the responsibility of the authors and do not necessarily represent the official views of NIAID or NIH. This work was also supported in part by the Bill & Melinda Gates Foundation grants INV-002143 (P.J.B.) and INV-036842 (M.S.S.).

Under the grant conditions of the Foundation, a Creative Commons Attribution 4.0 Generic License has already been assigned to the Author Accepted Manuscript version that might arise from this submission.

## Author contributions

Conceptualization: K.A.D., H.B.G., P.J.B.; Methodology: K.A.D., Y.E.L., Z.Y., P.N.P.G., M.S.S.; Formal Analysis: K.A.D., A.P.W.; Investigation: K.A.D., Y.E.L., Z.Y., P.N.P.G., M.S.S.; Resources: M.S.S., P.J.B.; Data Curation: K.A.D., Z.Y.; Writing - Original Draft: K.A.D., P.J.B.; Writing - Review & Editing: K.A.D., H.B.G., A.P.W., M.S.S., P.J.B.; Visualization: K.A.D.; Supervision: K.A.D., H.B.G., P.J.B.; Project Administration: K.A.D., P.J.B.; Funding Acquisition: M.S.S., P.J.B.

## Data Availability

The cryo-EM maps and atomic structures have been deposited in the Protein Data Bank (PDB) and Electron Microscopy Data Bank (EMDB) under accession codes 9EHL and EMD-48059 for the structure of IOMAmin5–BG505–10-1074 class I, and under accession codes 9EHM and EMD-48060 for the structure of IOMAmin5–BG505–10-1074 class II.

