## Supplemental Information for "Mapping essential somatic hypermutations in a CD4-binding site bNAb informs HIV-1 vaccine design"

**A**

```
                             5    10   15   20   25   30   35   40   45   50 52A55   60   65   70   75   80 82A  85   90     95  100          105  110
                                                        CDRH1                   CDRH2                                                  CDRH3
VH1-2*02            QVQLVQSGAEVKKPGASVKVSCKASGYTFTGYYMHWVRQAPGQGLEWMGWINPNSGGTNYAQKFQGRVTMTRDTSISTAYMELSRLRSDDTAVYYCAR----------------------------
IOMA-iGL-HC         ..................................................................................................EMFDSSADWSPWRGMVAWGQGTLVTVSS
IOMA-HC             E....E...Q........T...T....K....H.........R..........FR.AVK.P.N.R...S......MEIF.......T...........EMFDSSADWSPWRGMVAWGQGTLVTVSS
IOMA-HC-exFACE      ..................T........K....H....................FR.AVK.P.N.R...S......ME.........T...R.......EMFDSSADWSPWRGMVAWGQGTLVTVSS
IOMA-HC-exFACEmin   .........Q........T...T....K....H....................FR.AVK.P.N.R...S......MEIF...................EMFDSSADWSPWRGMVAWGQGTLVTVSS
IOMA-HCmin3         .........Q........T...T.........H....................FR.AVK.P...R...S......MEIF...................EMFDSSADWSPWRGMVAWGQGTLVTVSS
IOMA-HCmin4         .........Q........T...T.........H....................FR.AVK.....R...S.......EIF...................EMFDSSADWSPWRGMVAWGQGTLVTVSS
IOMA-HCmin5         .........Q........T...T.........H....................FR.AVK.....R...S.......EIF...................EMFDSSADWSPWRGMVAWGQGTLVTVSS
IOMA-HC-DDE         E....E...Q........T...T....K....H.........R..........FR.AVK.P.N.R...S......MEIF.......T...........EMFDSDDEWSPWRGMVAWGQGTLVTVSS

                             5    10   15   20   25 27A  30   35   40   45   50   55   60   65   70   75   80   85   90  95 100 105  110
                                                   CDRL1                   CDRL2                                        CDRL3
LV2-23*02           QSALTQPAS-VSGSPGQSITISCTGTSSDVGSYNLVSWYQQHPGKAPKLMIYEVSKRPSGVSNRFSGSKSGNTASLTISGLQAEDEADYYCCSYA-GSSTF-------------
IOMA-IGL-LC         ...............................................................................................-.VA.GGGTKLTVLGQPKA
IOMA-LC             .......................A.S.R...GFD...............I....N.....I.S...A...............E....H...Y...D.VA.GGGTKLTVLGQPKA
IOMA-LC-exFACE      .........................S.R...GFD................................A........................Y...D.VA.GGGTKLTVLGQPKA
IOMA-LC-exFACEmin   .........................S.R...GFD................................A........................Y...D.VA.GGGTKLTVLGQPKA
IOMA-LC-min3        .........................S.R...GFD................................A........................Y...D.VA.GGGTKLTVLGQPKA
IOMA-LC-min4        .........................S.R...G.D................................A........................Y...D.VA.GGGTKLTVLGQPKA
IOMA-LC-min5        .........................S.R...GFD................................A........................Y...D.VA.GGGTKLTVLGQPKA
IOMA-LC-5aa-CDRL3   .......................A.S.R...GFD...............I....N.....I.S...A...............E....H...Y..D---A.GGGTKLTVLGQPKA
IOMA-LC-9aa-CDRL3   .......................A.S.R...GFD...............I....N.....I.S...A...............E....H...Y...D.VA.GGGTKLTVLGQPKA
```

**B**

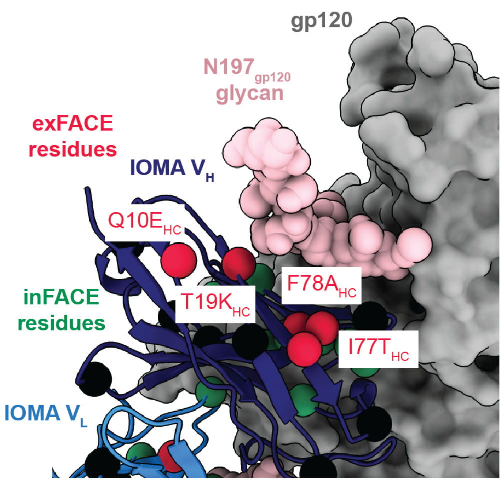


**C**

| | | # of SHM Subsitutions | | Percent mutation | |
|---|---|---|---|---|---|
| | | $V_H$ | $V_L$ | $V_H$ % | $V_L$ % |
| IOMA-class | IOMA | 22 | 15 | 22 | 14 |
| | IOMA exFACEmin | 18 | 8 | 18 | 8 |
| | IOMAmin3 | 16 | 8 | 16 | 8 |
| | IOMAmin4 | 14 | 7 | 14 | 7 |
| | IOMAmin5 | 14 | 8 | 14 | 8 |
| | ACS101 | 27 | 18 | 27 | 19 |
| | ACS102 | 35 | 16 | 35 | 17 |
| | ACS103 | 24 | 20 | 24 | 21 |
| VRC01-class | VRC01 | 41 | 23 | 41 | 25 |
| | minVRC01 | 14 | 7 | 14 | 8 |
| | 12a21 | 31 | 20 | 31 | 22 |
| | 12a21min | 17 | 8 | 17 | 9 |
| | BG24 | 23 | 19 | 23 | 20 |
| | PCIN63-71I1a | 21 | 17 | 21 | 18 |
| | N6 | 41 | 30 | 41 | 32 |
| | 3BNC117 | 34 | 23 | 34 | 24 |
| | NIH45-46 | 44 | 27 | 39 | 25 |
| | VRC03 | 40 | 36 | 40 | 38 |

**D**

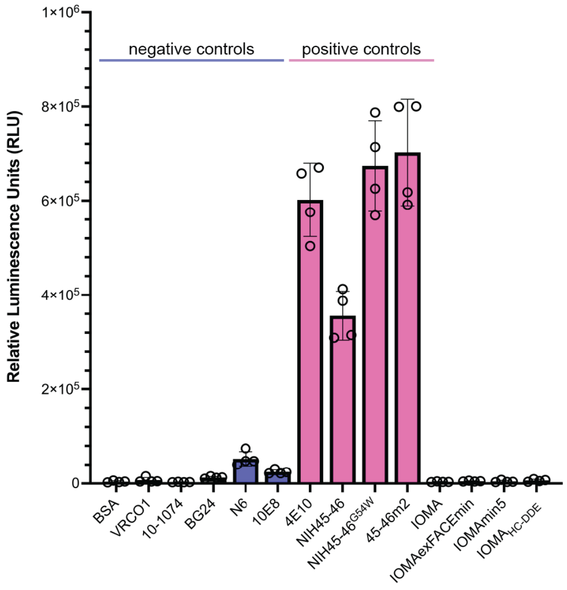

**Figure S1. Sequence alignment and characterization of IOMAmin antibodies, related to Figure 1.**

(A) Sequence alignment of IOMA iGL, mature, and mutant VH/VL sequences with CDR regions (as defined by Kabat numbering) highlighted in yellow. (B) Structural representation of IOMAexFACEmin Fab bound to gp120 with exFACE and inFACE residues shown as red and green spheres, respectively (PDB 5T3X). Remaining SHMs are shown as black spheres. exFACE mutations in closest proximity to the N197gp120 glycan are labeled. (C) Summary of the number of SHM amino acid substitutions and percent mutation for IOMA-class and VRC01-class bNAbs. (D) Results of a baculovirus polyreactivity ELISA-based assay[34] evaluating non-specific binding of bovine serum albumin (BSA), a panel of control HIV-1 IgG bNAbs, and IOMA mutant IgG bNAbs

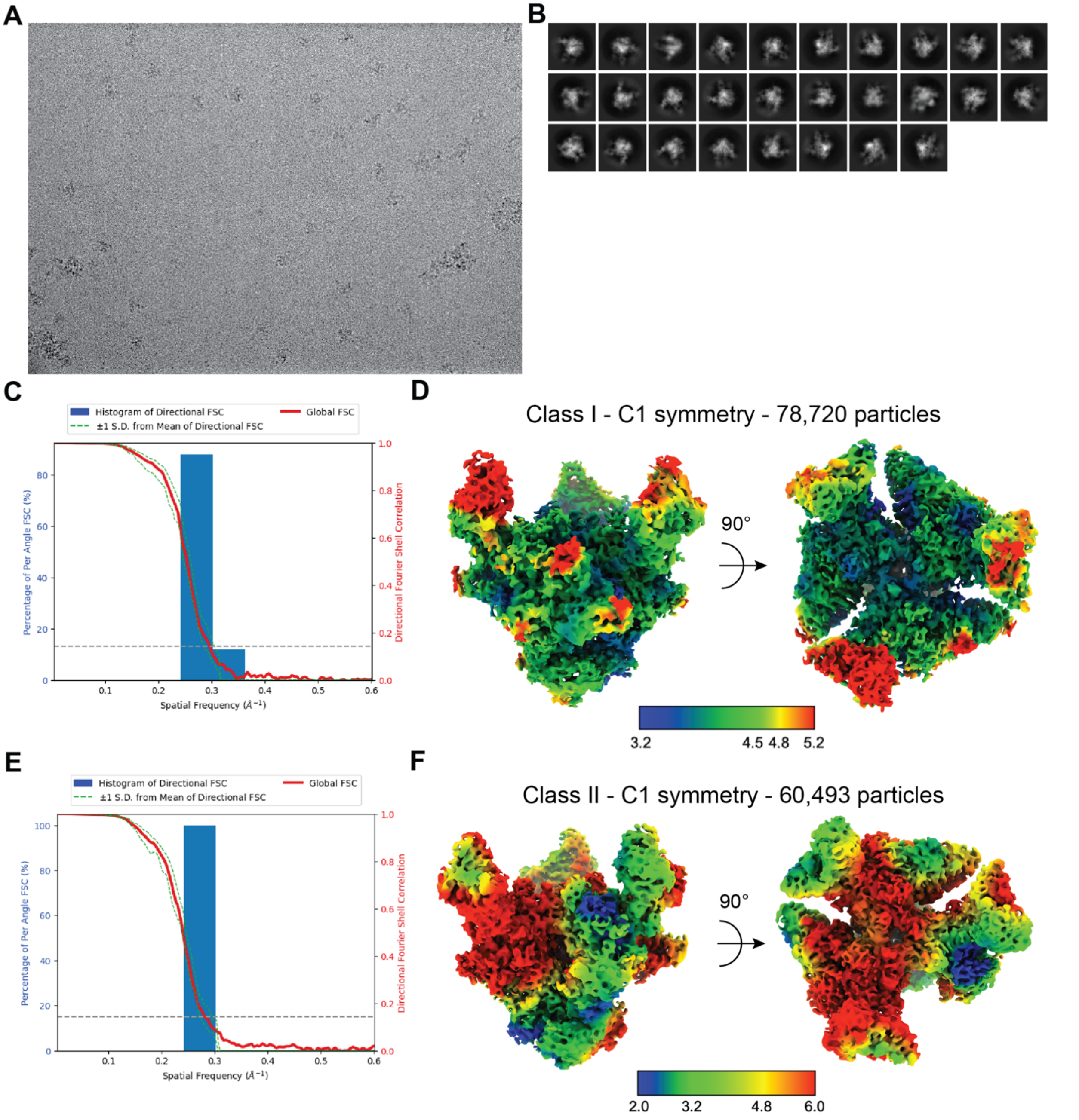


**Figure S2. Cryo-EM data processing and validation for IOMAmin5-BG505-10-1074 complexes, related to Figure 3.**

(A) Representative micrograph and (B) cryo-EM 2D class averages for the IOMAmin5-BG505-10-1074 cryo-EM structures. For this dataset, two classes were resolved: class I with three IOMAmin5 Fabs bound to BG505 and class II with two IOMAmin5 Fabs bound to BG505. (C) Gold-standard Fourier shell correlation (FSC) plot and (D) local resolution map for IOMAmin5-BG505-10-1074 class I. (E) Gold-standard FSC plot and (F) local resolution map for IOMAmin5-BG505-10-1074 class II.
