## Supplemental Table 1 for "Mapping essential somatic hypermutations in a CD4-binding site bNAb informs HIV-1 vaccine design"

### Table S1: Cryo-EM data collection, refinement and validation statistics

| | **IOMAmin5-10-1074-BG505 Class I** (EMDB-48059) (PDB 9EHL) | **IOMAmin5-10-1074-BG505 Class II** (EMDB-48060) (PDB 9EHM) |
|---|---|---|
| **Data Collection and Processing** | | |
| Microscope | Titan Krios | Titan Krios |
| Camera | Gatan K3 | Gatan K3 |
| Magnification | 105,000 | 105,000 |
| Voltage (keV) | 300 | 300 |
| Exposure (e-/Å2) | 60 | 60 |
| Pixel size (Å) | 0.4327 | 0.4327 |
| Defocus Range (um) | -1 to -3 | -1 to -3 |
| Initial Particle Image (no.) | 860,116 | 860,116 |
| Final Particle Image (no.) | 78,720 | 60,493 |
| Symmetry Imposed | C1 | C1 |
| Map Resolution (Å) | 3.9 | 4.2 |
| FSC Threshold | 0.143 | 0.143 |
| **Refinement** | | |
| Initial Model Used | PDB 5T3X | PDB 5T3X |
| Model Resolution (Å) | 3.9 | 4.2 |
| FSC Threshold | 0.143 | 0.143 |
| Model composition | | |
| Non-hydrogen atoms | 26,146 | 23,943 |
| Protein residues | 3,165 | 2,898 |
| Ligands | 113 | 99 |
| Average B-factors (Å2) | | |
| Protein | 95 | 104 |
| Ligands | 80 | 89 |
| R.m.s. deviations | | |
| Bond length (Å) | 0.006 | 0.007 |
| Bond angles (º) | 1.1 | 1.1 |
| Validation | | |
| MolProbity score | 2.0 | 2.2 |
| Clashscore | 12.4 | 18.7 |
| Rotamer outliers | 0.74 | 0.72 |
| Ramachandran plot | | |
| Favored (%) | 93.2 | 93.6 |
| Allowed (%) | 6.8 | 6.4 |
| Outliers (%) | 0 | 0 |
